# RNA structure conservation in plastids across plant evolution

**DOI:** 10.1101/2025.08.06.668526

**Authors:** Dolly Mehta, Chen Xiao, Jingmin Hua, Rodrigo S. Reis

## Abstract

Plastid genomes are deeply evolutionary conserved. RNA structures within the primary or mature transcript play central role in plastid regulation of RNA processing, stability, and translation. However, the identity and conservation of RNA structures selected in plastid’s evolution are still largely elusive. Here, we developed a stringent, covariation-based pipeline that perform an unbiased screen for conserved RNA secondary structures across entire plastid genomes. We analysed ∼14,000 plastid genomes and identified a repertoire of 57 high-confidence conserved structures. We recovered known functional classes, e.g., 16S rRNA, tRNA, group II intron, and 3′end stem-loop, evidencing that our genome-wide analysis is reliable. We further uncovered novel putative *cis*-acting structures within the UTRs and introns of key photosynthetic genes, including *psbN*, *clpP*, and *atpF*, as well as putative *trans*-acting antisense RNAs to *petB* and *psbT*, suggesting uncharacterized elements with major regulatory function. Experimental *in vivo* RNA probing demonstrated that nearly half of the conserved structures adopt the predicted conformation in Arabidopsis plastid. Our comprehensive, yet stringent atlas of conserved plastid RNA structures provides the foundations for new regulatory discoveries in plastid biology.

## INTRODUCTION

Plastids evolved from endosymbiotic cyanobacteria likely ∼1.5 billion years ago (Bya) in a common ancestor shared by land plants and algae (Parfrey et al., 2011; Shih and Matzke, 2013; Archibald, 2015). In land plants, chloroplast genomes are conserved in structure and organisation, typically ∼150 kb circular quadripartite DNA sequence with two copies of an inverted repeat (IR) separated by a large and small single copy (LSC and SSC) regions, and 100-120 genes, mostly encoding subunits of the photosystems, electron transport chain, and gene expression machinery (Tonti Filippini et al., 2017). Substantial differences are, however, observed in some plant clades, from the absence of “essential” genes in grasses (Guisinger et al., 2010) and lack of an IR copy in Fabeae and related legumes (Palmer and Thompson, 1982; Wu et al., 2011), to extreme cases where the chloroplast lost its photosynthetic function, such as in parasitic plants (Wicke et al., 2016). The latter comprises the smallest plastid genomes, reaching as short as ∼11 kb, while few genomes are larger than 200 kb (Tonti Filippini et al., 2017). Despite some variations, plastid genome in land plants is far more conserved than mitochondrial and nuclear genomes, possibly because of a lower mutation rate in plastids (Smith, 2015), indicating stronger evolutionary pressure on a likely optimal sequence composition shared among land plants.

Plants have evolved to rely on plastids for various functions beyond photosynthesis, including development and environmental adaptation (Bobik and Burch-Smith, 2015). Compared with the cytoplasm, plastid provides cellular compartments with unique properties, such as distinct pHs (i.e., ∼8.0 and ∼5.5 in stroma and thylakoid lumen, respectively), reducing environment, and high levels of O_2_, CO_2_, Mg^2+^, and ATP (Kirchhoff, 2019). Many metabolites are primarily or exclusively produced in plastids, including starch, chlorophylls, carotenoids, glutathione, fatty acids, and some amino acids and hormones (Neuhaus and Emes, 2000). In fact, as opposed to mitochondria, the same plastid genome (ptDNA) is shared by diverse and specialised organelles, such as chloroplast and non-photosynthetic amyloplast (storage of starch), elaioplast (fatty acid), chromoplast (pigments), proteinoplast (protein), gerontoplast (aging chloroplast), and etioplast (non-photosynthetic chloroplast in dark-grown plants) (Neuhaus and Emes, 2000). Nuclear and plastid gene expression determine such specialisation—a process permissive to interconversion in response to environmental cues, developmental stage, and tissue type (Choi et al., 2021). For instance, chloroplast to chromoplast conversion is responsible for the green to red transition in tomato fruit during ripening (Powell et al., 2012), while potato tubers exposed to light have their amyloplasts converted into chloroplasts in peripheral cell layers, causing tuber greening via accumulation of chlorophyll (Tanios et al., 2018). Among the different plastids, chloroplast has a broader role in plant development and adaptation, not only for the importance of photosynthesis, but also for the conserved retrograde signalling pathways, by which chloroplast regulate nuclear gene expression via selective release of specific signals in response to cellular and environmental cues (Chan et al., 2016; Richter et al., 2023; Lee and Kim, 2024).

Evolution of plastid gene expression regulation is central to plant biology. Indeed, there has been a wealth of conservation studies on plastid genes and proteins (Martín and Sabater, 2010; Wicke et al., 2011; Guo et al., 2017), revealing, for instance, deeply conserved prokaryotic-type RNA polymerase (Vergara-Cruces et al., 2024; Wu et al., 2024) and ribosome (Bieri et al., 2017; Graf et al., 2017) similar to those in cyanobacteria. Most plastid genes are organised as operons, and chloroplast has evolved specialised post-transcriptional RNA processing to ensure transcript diversity and proper stability and translation of mature transcripts, such as RNA editing, splicing, 5′ and 3′ trimming, and intercistronic cleavage (Stern et al., 2010). Regulation of RNA processing, stability, and translation often require the presence of RNA structure within the primary or mature transcript (Zoschke and Bock, 2018). However, whether and how these regulatory elements are conserved are still poorly understood, and most well-characterized examples are limited to common cases for RNA structure, such as tRNA (Lohan and Wolfe, 1998), rRNA (Tiller and Bock, 2014), and self-splicing intron (Vogel, 2002; Stern et al., 2010), or functional analysis of RNA structure in one or few plant species (Zhou et al., 2007; Prikryl et al., 2011; Scharff et al., 2011; Hammani et al., 2012; Scharff et al., 2017). For instance, presence of stable stem-loop structure at the 3′ end of many primary transcripts is well-known, as this is a critical motif for precise 3′ end processing and RNA stability (Stern and Gruissem, 1987), while much less is understood about their conservation, including nucleotide pairing covariation and stem-loop length.

In this work, we present an unbiased genome-wide identification of conserved RNA secondary structures in plastids. We developed a computational approach for efficient, robust, and stringent covariation analysis using multiple sequence alignments (MSAs) produced with sliding windows covering the entire plastid genome of thousands of plant species. We compared pairing probability matrices for each sequence in each MSA to identify the shared secondary structure in the alignments, i.e., no a priori structure was used (e.g., minimal free energy structure). We then retained only MSAs associated with structures ranked above stringent criteria for accepting motifs. We further applied R-scape (Rivas et al., 2016) to retain RNA motifs with well-supported evidence for conservation. We could identify high-confidence 57 RNA motifs conserved across ∼14k plastid genomes, including known functional motifs (e.g., 16S rRNA, tRNA, group II intron, and 3′ end stem-loop) and novel uncharacterized motifs, such as within the UTR of *psbI*, *rps4*, *rps18*, *clpP*, *rpoB*, and *psbN*, and intronic region of *atpF* and *rpl16*. In addition to *cis*-acting RNA motifs, we identified conserved structure in antisense RNAs to *petB*, *petD*, *rps16*, and *psbT*-*psbB*. In addition, experimental RNA probing confirmed that most conserved structures are formed *in vivo*. Our work establishes a framework for reliable identification of conserved RNA structures within large genomic datasets and unveils novel putative regulatory elements in plastid RNA maturation, stability and/or translation.

## RESULTS AND DISCUSSIONS

### Genome-wide identification of conserved RNA secondary structures in plastids

Identification of conserved RNA structure requires multiple sequence alignment, typically, built using pre-specified target sequences (Stav et al., 2019; Gao et al., 2021; Brewer et al., 2021). Tools for screening of entire genomes, such as RNAz (Washietl et al., 2005), are based on calculation of thermodynamic stability (mainly favours minimum free energy structures), which commonly leads to spurious predictions and are, thus, not suitable for identification of motifs (Will et al., 2007; Ziesel and Jabbari, 2024). A powerful approach is to leverage the existing repertoire of conserved RNA structures available from the RNA family (Rfam) database (Ontiveros-Palacios et al., 2025). For this, we used the *cmsearch* program of Infernal (Nawrocki and Eddy, 2013) to screen the ∼14k plastid genomes available from NCBI. We identified conserved motifs for tRNAs, rRNAs, group I and group II introns, and the isrR-antisense RNA (Supplementary Table 1). The latter was found in nearly all plastid genomes and mapped to an antisense region within the *psbC* gene. In iron-deficient cyanobacteria, IsiA (iron stress-induced protein A) forms a giant ring structure around photosystem I, a process post-transcriptionally regulated by the expression of an internal IsiA antisense noncoding RNA, IsrR (iron stress-repressed RNA) (Dühring et al., 2006). IsrR-antisense RNA is conserved across photosynthetic species, suggesting that this noncoding RNA is a central, yet uncharacterized regulator of photosynthesis in most plant species. Therefore, search using the Rfam database proved valuable, but limited to the identification of a single putative novel motif in plastids. We, thus, developed a pipeline for automated, fast, stringent, and reliable screening of thousands of genomes for the identification of conserved RNA motifs. This was applied to plastid genomes, albeit it is in principle applicable to any set of genomes.

Seed evolution ∼370 Mya was a pivotal innovation in plants, with seed plants (i.e., gymnosperms and angiosperms) accounting for over 90% of living species (Baroux and Grossniklaus, 2019). Their current success involved diversification of photosynthetic strategies to support environmental adaptation, such as the evolution of C3, C4, and CAM photosynthesis (Edwards, 2019). We, thus, obtained 3823 manually curated plastid genomes from seed plants (Singh et al., 2020), and extracted sliding window sequences of 250 nucleotide (nt) in length, with 75 nt of overlap, covering the entire genomes (∼5.6M windows) (Fig. 1a). We removed redundant windows and produced clusters with a minimum of 10 windows and 50% sequence identity using MMSeqs2 (Steinegger and Söding, 2017), resulting in >10,000 clusters. A pre-screening was performed with the fast-performing RNALalifold (Lorenz et al., 2011), which calculates locally stable secondary structures, to identify clusters with structures sharing at least 2 base pairs. This resulted in 3136 clusters, for which each sequence was refolded and realigned using the probabilistic mode of LocARNA (Will et al., 2012), producing 1827 clusters where sequences share secondary structure (Fig. 1b). For these clusters, we evaluated whether each sequence within an alignment adopts the consensus structure to produce clean alignment with only the sequences forming the predicted structure. This resulted in 667 clusters (out of 1827) with a minimum of 7 non-redundant sequences. We used these cleaned alignments as seeds to find homolog sequences using Infernal (Nawrocki and Eddy, 2013), which were then processed to obtain alignments including the new hits. Sequences that were non-redundant and formed the predicted structures were retained, thus producing improved alignments with better performance as seed. We then evaluated these alignments for covariation using R-scape (Rivas et al., 2016) to identify conserved RNA structures beyond phylogenetic expectations. R-scape identified 145 clusters (out of 667) with at least 2% base pairs showing statistically significant covariation (Fig. 1b). Alignments obtained with LocARNA are often biased towards positions containing structure, hence, we further evaluated the alignments using nhmmer (Wheeler and Eddy, 2013) and CaCoFold (Rivas, 2020) to realign the sequences without structure information, evaluate the LocARNA consensus structure, respectively, and then retained only the alignments that support the corresponding predicted structure by LocARNA. Out of 145 clusters, 76 passed this evaluation, thus, corresponding to RNA motifs with covariation for which we established high confidence predicted structure and high alignment reliability. We then used the multiple sequence alignment for each conserved RNA motif as covariance models to search for structure homologs across ∼14k plastid genomes, from non-seed plants to evolutionarily distant unicellular photosynthetic organisms (Fig. 1a). We reasoned that conserved structure displaying consistent context (e.g., similar position in a gene) across plastid genomes is more likely to be a true conserved regulatory element. Therefore, we mapped the genomic context for each motif and identified 57 motifs (out of 76) with regulatory potential, while the remaining 19 motifs were intergenic without readily identifiable putative function. We thus focused on those 57 conserved motifs with regulatory potential, hereafter, conserved RNA motifs in plastids (Fig. 1a-b). Most of these motifs (∼85%) are associated with noncoding (rRNA, tRNAs, and antisense RNA) or coding (UTR, exon-intron junctions, and CDS) genes, while genic and functional association for ∼15% is unclear (Fig. 1c).

**Figure 1.**
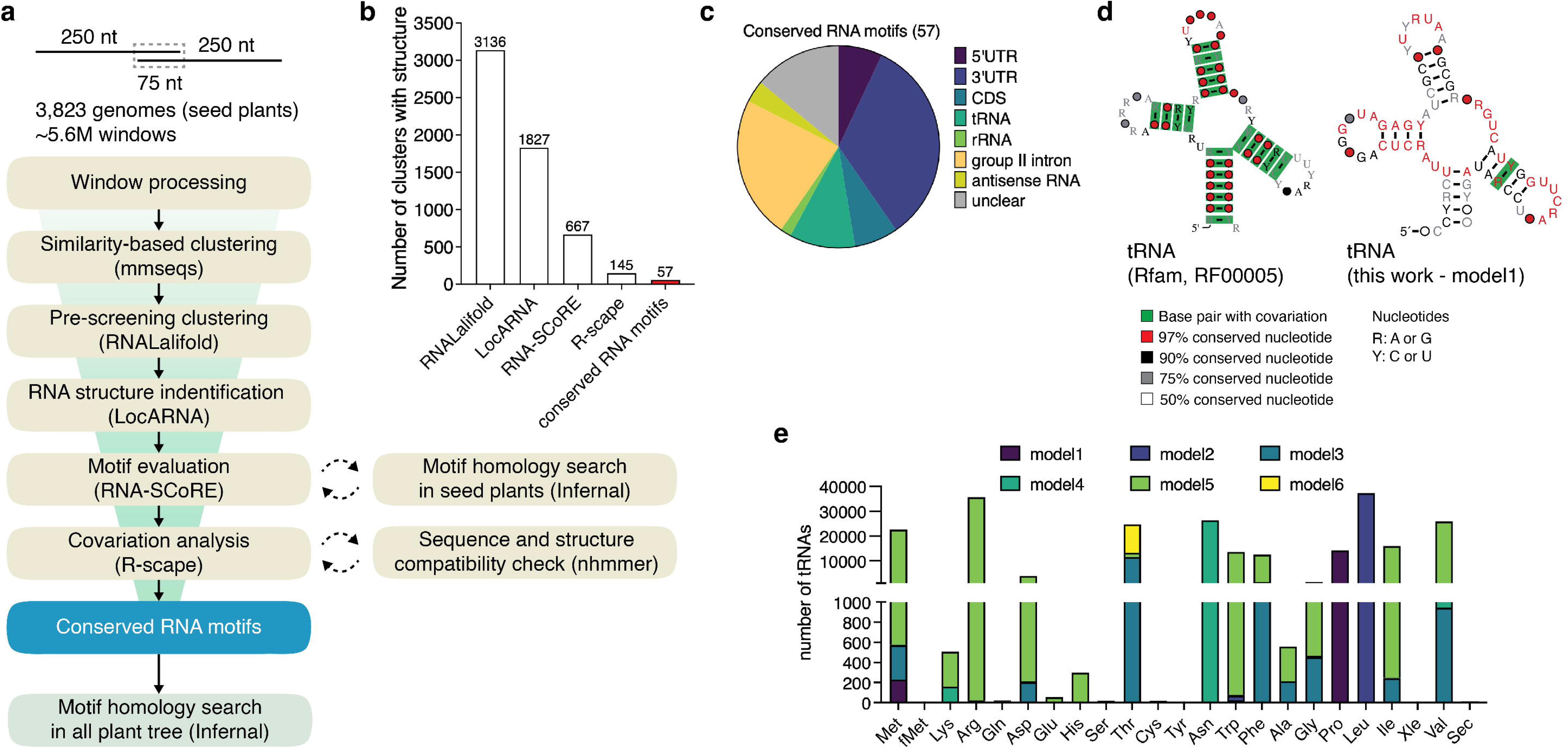
Identification of conserved RNA structure in plastid genomes. **(a)** Pipeline to predict and evaluate conservation of secondary structure in plastid genomes. **(b)** Bar plot showing the number of clusters at each pipeline step, resulting in the identification of 57 high confidence conserved RNA motifs (red bar). **(c)** Pie chart showing the genomic context for the conserved RNA motifs. **(d)** Consensus structure for tRNA obtained from Rfam database (left) and this work (right). Covarying base pairs (e-value ≤ 0.05) are highlighted in green, as shown in the legend. **(e)** Stacked bar plot showing the number of tRNAs identified across 14034 plastid genomes with six tRNA models.

tRNA function is based on folding into specific conformation, and it is, therefore, expected to display covariation in our analysis. We obtained six models mapping to annotated plastid tRNAs, all of which also identified using the Rfam model for tRNA (RF00005) (Fig. 1d, Supplementary Fig. 1a), i.e., 100% positive predictive value (PPV). The Rfam model displays covariation in all base pairs, while fewer were identified in our tRNA models, likely explained by higher sequence identity among tRNAs in plastids, as compared to phylogenetically distant genomes used in the Rfam model (Lowe and Eddy, 1997). While two of our tRNA models (model1 and model4) mapped to different types of tRNAs, the other four models showed amino acid specificity (Fig. 1e), indicating that our pipeline captures functionally related structures with relatively subtle differences that are important for function. We also identified a conserved 16S rRNA structure motif mapping to bacterial helix 8, matching the Rfam model for 16S rRNA RF00177 (Supplementary Fig. 1b and Supplementary Table 2). Our models identified tRNAs and 16S rRNA helix in plastids across the entire plant phylogenetic tree with high sensitivity, i.e. covering over 99% of plant species compared to Rfam models (Fig. 2), evidencing that our pipeline for the identification of conserved structures across genomes is effective and reliable. The conserved structures identified here are already available in Rfam, the main database of RNA families, under the accessions listed in Supplementary Table 2.

**Figure 2.**
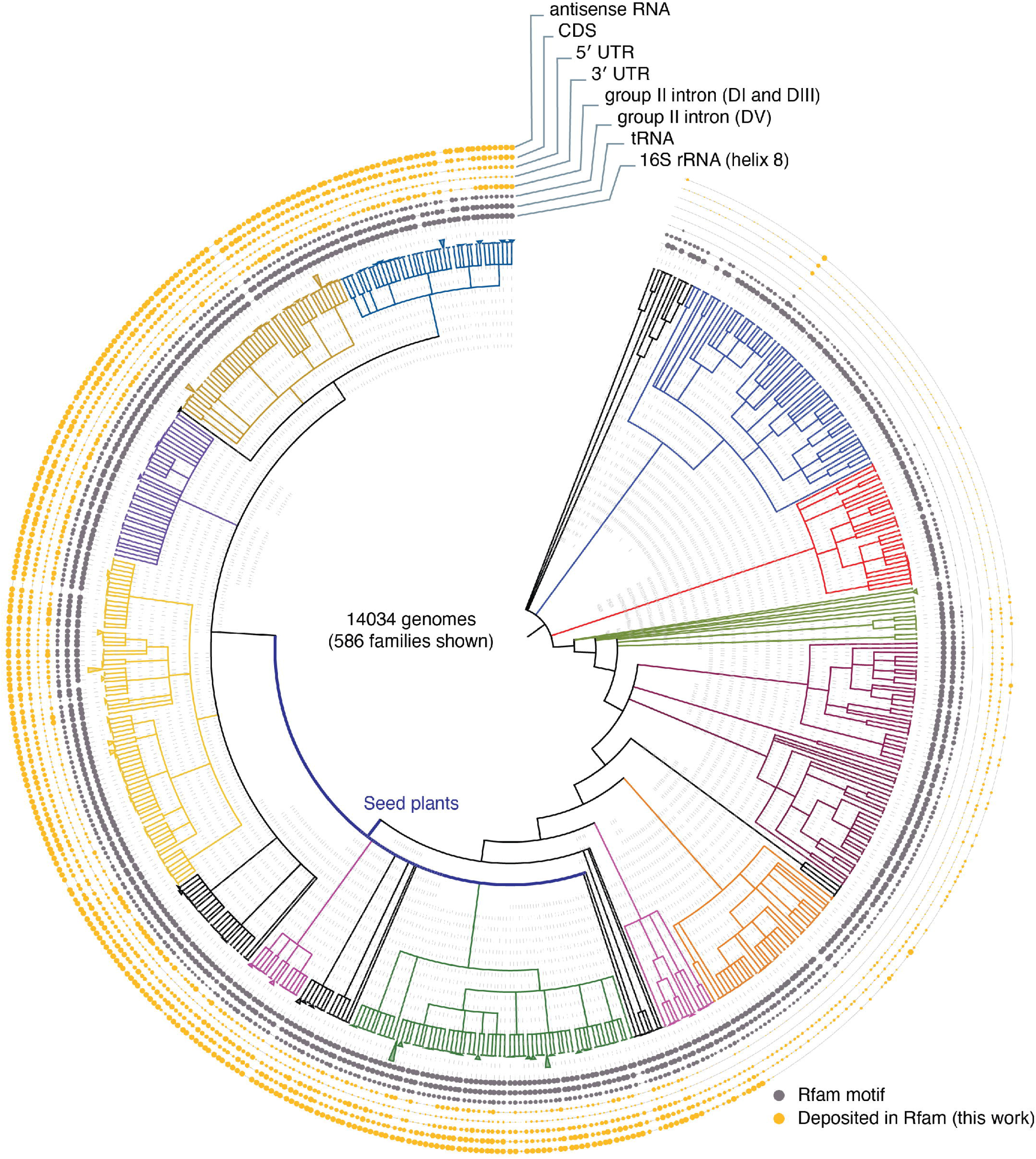
Distribution of conserved RNA motifs in the phylogenetic tree of Viridiplantae. Circle size (outer layer) represents the percentage distribution of RNA motif in given plant clade (coloured branches). Grey circle represents RNA motifs available in Rfam (i.e., known motifs) and yellow represents novel conserved motifs. RNA motifs are grouped based on genomic context and cellular functions. Seed plants are highlighted in blue.

### Experimental validation of *in vivo* folding for conserved RNA structures

Next, to assess whether the conserved structures fold into the predicted conformation *in vivo* under standard plant growth conditions, we used dimethyl sulfate (DMS) combined with mutational profiling (MaP) to chemically probe RNA structures inside plastids (Zubradt et al., 2017). DMS preferentially modifies unpaired adenine and cytosine, in which the mutation rate induced by DMS modification reveals the nucleotide structural state. Our DMS MaP data for the plant model *Arabidopsis thaliana* Col-0 (hereafter, Arabidopsis) had sufficient coverage for the sequence of nearly all conserved structures, i.e., 55 out 57 (Supplementary Table 3). We then evaluated the agreement between each conserved structure and the corresponding reactivity profile. For that, we calculated the area under the receiver operating characteristic curve (AUROC), which is the area underlying the curve described by the set of false and true positive rate (FPR-TPR) value pairs at each value of reactivity threshold (0 to 1), i.e., a high AUROC value indicates better agreement between the structure and reactivity profile. We obtained an AUROC ≥ 0.50 for 44 structures, out of which, after manual curation to avoid agreement misassignment, we confirmed reactivity support for 25 (labelled in green in the Supplementary Table 3), while the remaining structures were either unclear or ambiguous (12; labelled in yellow) or not supported (7; labelled in red). In total, only 18 out of 57 structures were not supported by DMS MaP data. Among the 25 conserved structures experimentally validated for their *in vivo* folding, we recovered tRNA structures showing low reactivities in the base-pairs and showing highly reactive unpaired nucleotides (Supplementary Fig 1c), demonstrating successful *in vivo* DMS probing. Furthermore, we obtained probing data for several other motifs identified in this study (described below).

### Conservation in motifs within group II intron

Plastid genome has an origin in cyanobacterium and has retained self-splicing introns within a subset of their genes, albeit plastids evolved to require protein cofactors (mostly nuclear-encoded) necessary for efficient and proper splicing (Small et al., 2023). They belong to two main classes of self-splicing RNAs, i.e., group I and II introns, and their catalytic activity is intrinsically linked to their ability to fold into specific and highly conserved conformation— required for both the ribozyme activity and RNA interaction with protein cofactors (Small et al., 2023). In land plants, ∼20 group II introns and one group I intron have been reported in plastid genomes (Asakura and Barkan, 2007). Here, we identified conserved structures in group II introns mapping to NADH dehydrogenase (*ndhA* and *ndhB*), ribosome subunit (*rpl16* and *rps12*), photosystem I assembly (*ycf3*), cytochrome B6f complex (*petB* and *petD*), ATP synthase (*atpF*) and tRNA (*trnK* and trnG) (Supplementary Fig. 2a).

Group II intron is defined by a characteristic and conserved secondary structure, typically consisting of six major stem-loop domains, designated D1, D2, D3, D4, D5, and D6 (Fig. 3a). These domains radiate from a central core region (multiloop junction), forming an arrangement that is essential for bringing distant sequence elements, including the 5’ and 3’ splice sites and the branch-point nucleotide, into the precise spatial orientation required for splicing (Lambowitz and Zimmerly, 2011). D5, a key catalytic domain, is the most conserved domain (structurally and in sequence), typically forms a short stem-loop structure (∼34 base pairs), and contains a conserved catalytic nucleotide triad (partially forming the AC bulge; Fig. 3a) and a GNRA tetraloop involved in D5 stabilisation and interaction with divalent metal ions and other regions of the intron. In fact, D5 shares the same structural conservation between group II intron (Fig. 3b) and U6 spliceosomal RNA, indicating an evolutionary link between self-spliced group II introns and splicing in eukaryotes (Shukla and Padgett, 2002).

**Figure 3.**
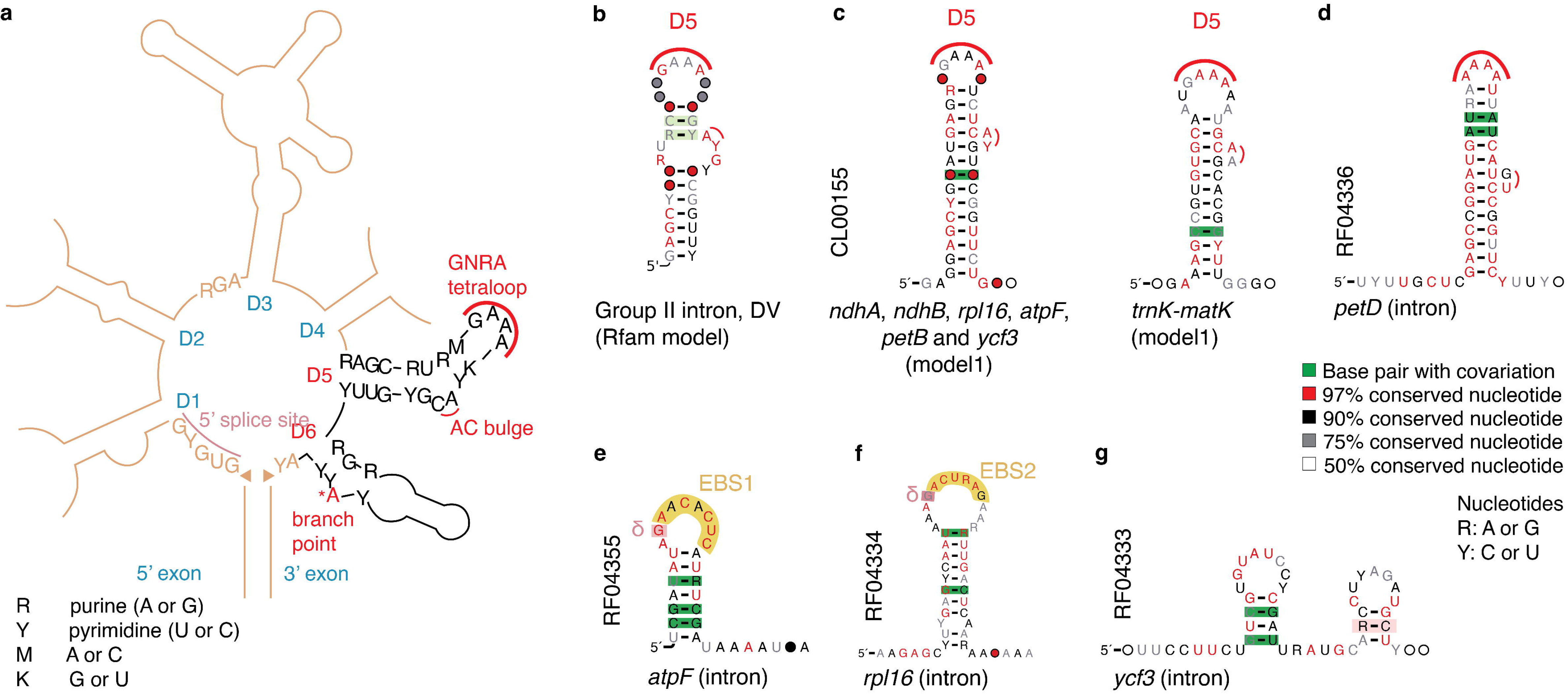
Conserved RNA motifs in group-II intron. **(a)** Simplified model for group II intron showing the typical six domains (D1-6) and key nucleotides involved in splicing. **(b)** Consensus representative structure model for D5 available in Rfam (Clan CL00102). Highly conserved GNRA loop and AY bulge highlighted with red line. **(c)** Consensus structure for two motifs identified in this work mapping to intronic regions. Conserved D5 with GNRA loop and AY bulge, highlighted with red line, frequently observed in *ndhA*, *ndhB*, *petB*, *atpF*, and *ycf3* introns, while AA bulge was found in *trnK* intron. **(d)** Consensus structure for motif identified in *petD* intron resembling D5, except for variation in loop and bulge sequence identity. **(e)** Consensus motif for D1 in *atpF* involved in tertiary interactions. EBS1, exon-binding site 1. **(f)** Consensus structure for D1 in *rpl16* highlighting the conserved EBS2. **(g)** Consensus structure for D1 in *ycf3* identified based on sequence position. **(b, d-g)** Rfam accessions provided on the left side of the structures (vertical text).

The stem-loop domain 6 (D6), on the other hand, harbours an unpaired bulged nucleotide, typically an adenosine, that serves as branch point for lariat formation (Fig. 3a), in which the adenosine’s 2’-hydroxyl group initiates the first transesterification reaction of splicing.

Our analysis identified conserved D5 represented by five models mapping to *ndhA*, *ndhB*, *petB*, *atpF*, *rps12*, *rpl16*, *rpoC1*, and *ycf3* (Fig. 3c and Supplementary Fig. 2b), and another two different models mapping to D5 in *trnK* (Fig. 3c and Supplementary Fig. 2c). We also found a conserved motif in *petD* that resembles D5 (Fig. 3d); however, this motif diverges in the loop (AAAA (Borsch et al., 2009; Korotkova et al., 2009)), instead of canonical GNRA tetraloop) and within the catalytic nucleotide triad (GU, instead of AC bulge). Furthermore, these group II introns were validated for their *in vivo* folding in Arabidopsis plastids using DMS-MaP (Supplementary Fig. S2d). Probing data also revealed that the unpaired nucleotides in the terminal loop of *petD* D5, unlike others, had low reactivity to DMS, suggesting that *petD* D5 group II intron is likely in a different folding state such as involving tertiary interactions (e.g., ζ-ζ’ with D1) or protein interactions for regulatory and/or catalysis functions. *petD* has a single group II intron processed by similar proteins required for processing of *ndhA*, *ndhB*, *rpl16*, *ycf3*–int1, among others (Asakura et al., 2008), suggesting that the differences in D5 between *petD* and aforementioned transcripts might have a regulatory function, but group II intron in *petD* likely retained similar processing. In accordance, *petD* is transcribed as part of the *psbB* operon (*psbB*-*psbT*-*psbH*-*petB*-*petD*), which has two group II introns (*petB* and *petD*), and null mutation in *ALBINO EMBRYO AND SEEDLING* (*AES*) gene results in similar intron retention in *petB* and *petD* (An et al., 2023). All mapped genes are known to harbour group II intron (Wang et al., 2022), and our findings indicate strong structural conservation for D5 and regulatory function associated with a D5 variant.

We also identified a conserved structure in the group II intron domain D1 of *atpF*, *ycf3*, *rpl16*, and *clpP* and D3 of *trnK* (Fig. 3e-g and Supplementary Fig. 3a). Typically, D1 is the largest and most variable of the six domains, which is further organised in subdomains that include conserved sequence elements known as exon-binding site 1 (EBS1) and EBS2. EBSs base pair with corresponding complementary intron-binding site 1 (IBS1) and IBS2 in the upstream exon to ensure accurate 5’ splice site recognition. D1 is also critical for global stabilisation of the intron RNA via tertiary interactions (e.g., δ-δ’, a-a’, and b-b’) (Lambowitz and Zimmerly, 2011). The conserved motif identified here is the stem-loop containing EBS1 and the guanine ( δ) that makes tertiary interaction with the first 3′exon nucleotide (δ′) adjacent to D6 (Inaba-Hasegawa et al., 2021) (Fig. 3e). EBS1–IBS1 pairing has been shown to be essential for splicing, whereas EBS2–IBS2 pairing is necessary for efficient splicing but is not required (Inaba-Hasegawa et al., 2021). In the same work, the authors showed that δ-δ′ interaction is not involved in efficient splicing and likely does not contribute to the 3′ splice-site selection, indicating a mechanistic difference between self-splicing group II intron and protein-mediated group II intron splicing in higher plant chloroplasts. We also found a structure in the 5’ end of *rpl16* as part of the group II intron D1, albeit it includes EBS2 binding site as previously reported for Japanese black pine (*Pinus thunbergii*) (Kelchner, 2002), instead of EBS1 found in the *atpF* motif (Fig. 3f). Furthermore, *in vivo* folding of these *rpl16* and *atpF* group II intron D1 were experimentally validated using DMS-MaP (Supplementary Fig. 3b). We found another motif in the group II intron D1 in *ycf3* and *clpP* (Fig. 3g, Supplementary Fig. 3a) based on its sequence position within the intron, while its position within the structured D1 is unclear. Furthermore, we identified a motif in *trnK* likely part of the group II intron D3 (Supplementary Fig. 3e), as compared with a *trnK* intron sequence in *Nicotiana tabacum* and relatively mapped to D3 of group II in *trnK* intron (RNAcentral: URS0002349F48/4097) (Cannone et al., 2002). Altogether, we obtained conservation models for group II intron D5 additional to previously available Rfam model (deposited as part of a clan), and motif within D1 that have not been previously described in Rfam (Supplementary table 2).

### Conservation in 3′ termini processing motif

Plastid transcription is typically pervasive and termination is inefficient, producing transcripts that require processing (Stern and Gruissem, 1987). Transcript processing often involves the presence of an inverted repeat (IR) in 3′ UTR to evade 3′-to-5′ exonuclease degradation and define precise 3 ′ end (Stern and Gruissem, 1987). Although terminal IRs are well-documented functional motifs in plastids, their conservation in sequence and structure is less well understood. Here, we identified 13 motifs representing conserved terminal IR motifs associated with genes involved in photosystem II (*psbA*, *psbM*, and *psbJ*), cytochrome c biogenesis (*ccsA*), and NDH complex (*ndhD*) (Fig. 4a-b and Supplementary Fig. 4a-c). These conservation models are mainly associated with three genes and, given that they identify corresponding motif across the phylogenetic tree, the models were combined into three clans: *psbA*, *psbJ*, and *ccsA*-*ndhD* (Supplementary Fig. 4d). IR-dependent RNA maturation and stability have been described for *psbA* (Stern and Gruissem, 1987; Adams and Stern, 1990) and *ndhD* (Zandueta-Criado, 2004), both of which with similar IR sequences as those identified here. Structural predictions have identified IR in the 3′ UTR of *ccsA*, *psbJ*, and *psbM* in various plant species (Kim and Lee, 2005; Jo et al., 2019), also similar to *psbA* and *ndhD*. Conserved IR motifs displayed 19-24 base pairs in the main stem-loop, including a bulge in *psbA* and *psbJ*, while there were no unpaired nucleotides in the IR for *ccsA*, *psbJ*, and *psbM*. We further confirmed that the 3′ termini IRs adopt the predicted structures *in vivo* (Fig. 4c). We also found that *psbJ* only forms the larger stem-loop *in vivo*, hence the shorter stem-loop is likely dispensable in RNA maturation. Furthermore, there were other shorter stem-loops as part of the conserved motifs in *psbA* and *psbJ*, both upstream the IR; these showed no covariation, suggesting that they might participate in regulatory processes other than RNA maturation. For instance, we found that the short hairpin in *psbA*, located upstream to the stop codon and protein binding site (Memon et al., 1996; Kupsch et al., 2012), forms *in vivo* (Fig. 4c), suggesting a regulatory role in transcript stability or translation in response to temperature changes.

**Figure 4.**
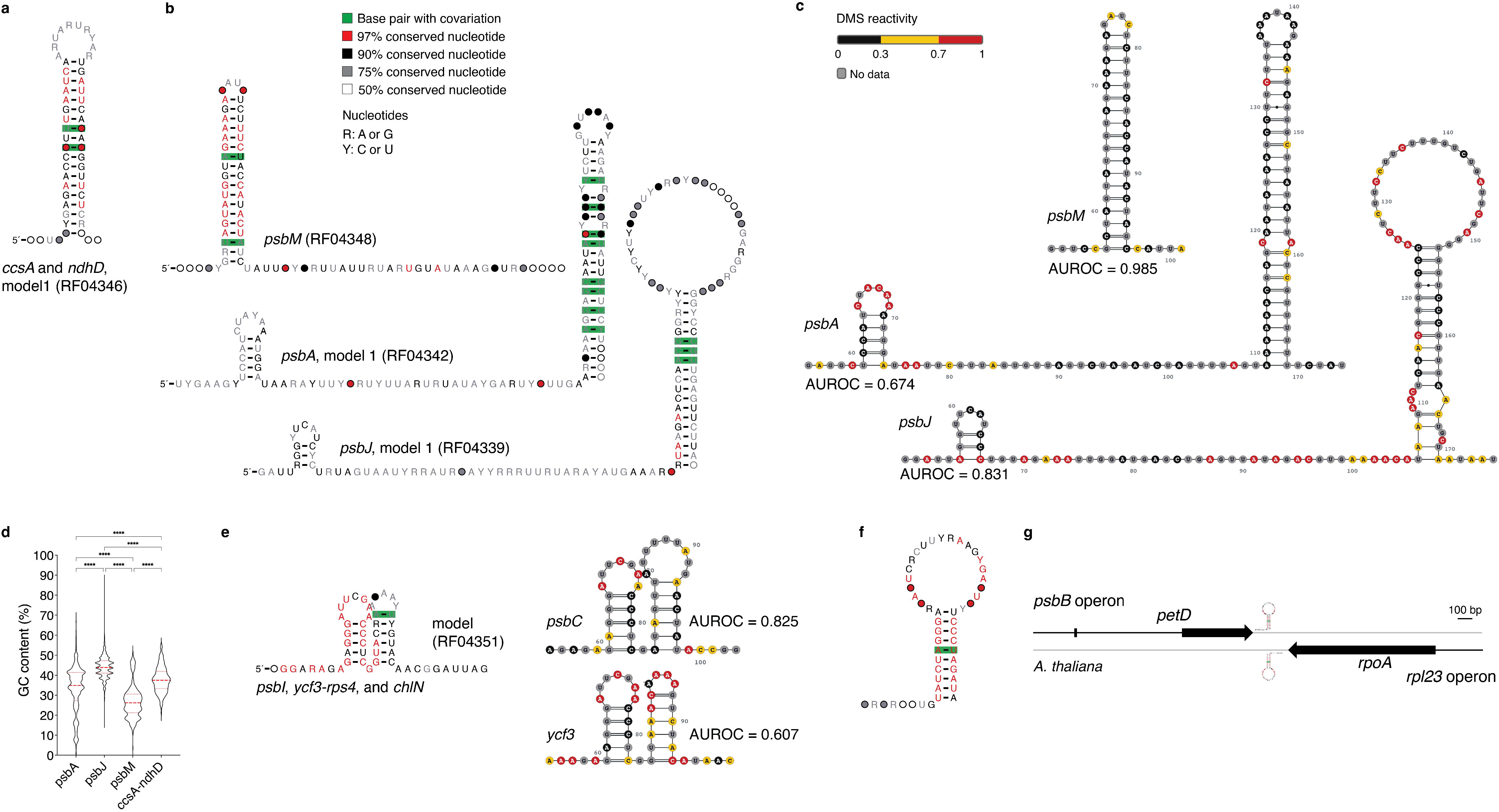
Conserved structure in 3’ terminal region. **(a-b)** Consensus structure for representative motifs involved in 3’ termini processing. **(c)** DMS reactivity for structures in **(b)** identified in *Arabidopsis thaliana*. Value <0.3 (black), 0.3-0.7 (yellow), and >0.7 (red) indicate low, medium, and high reactivity (i.e., unpairness), respectively. Area under the receiver operating characteristic curve (AUROC; 0 to 1) indicates the agreement between structure and reactivity profile, in which high AUROC values correspond to better agreement. **(d)** Violin plot for GC content percentage of sequences within each model shown in **(a)** (one-way ANOVA, p-value < 0.0001). **(e)** Consensus structure for a short tandem motif in the 3’ end of *psbI*, *ycf3-rps4*, and chlN (top) and corresponding DMS reactivity for Arabidopsis *psbC* (middle) and *ycf3* (bottom). **(f)** Consensus structure for a putative transcription terminator. **(g)** Genomic context of putative transcription terminator **(f)** in the *Arabidopsis thaliana* plastid.

We found the conserved motif in *psbM* across the phylogenetic tree, including in plastid of early flowering plant species such as *Amborella trichopoda*. There is evidence for widespread IR formation in 3′ends via small inversions in plastid genomes (Kim and Lee, 2005), which is expected to produce stems with perfect base pairing, while mismatches likely arise during evolutionary selection, as observed for stem-loop in plant miRNA precursors (Axtell and Bowman, 2008). We found that the *psbM* IR structure forms *in vivo* and displays the largest AUROC value in our dataset (0.985), suggesting a highly stable structure (Fig 4c). Stem-loop stability typically shows positive correlation with GC content (Becskei and Rahaman, 2022). Hence, we assessed GC content distribution in aligned sequences for *psbA*, *psbM*, *psbJ*, *ccsA*, and *ndhD* representative covariation models (Fig. 4d). We observed a broad range of median GC content, from 26% in *psbM* to 44% in *psbJ*, with relatively tight spread in the *psbJ* motif, suggesting that its higher GC content has been under evolutionary pressure. Selection of GC content in the *psbM* motif, on the other hand, appears weak and permissive to stem-loop with likely poor stability. It is, thus, possible that IRs display distinct stability with regulatory function, such as weaker evasion of 3′-to-5′ exonuclease degradation under conditions that destabilise IRs with lower GC (e.g., heat stress).

In addition to typically long terminal stem-loop, we identified a conserved terminal motif formed by two short stem-loops in tandem (Fig. 4e). This motif is found in the 3′ of *psbI*/*psbC* and *rps4* in most seed plants, but also in the 3′ of *ycf3* in some seed species and, in fewer cases, in the 3′ of chlN in evolutionarily distant species, such as single-cell green alga *Chlamydomonas reinhardtii*. This tandem terminal motif has been detected in *rps4* by 3’ RACE, where the authors showed evidence for protein interaction as a major mechanism to define 3’ transcript termini in plastids (Zhelyazkova et al., 2012b), albeit no detailed characterisation was performed for this particular motif. In Arabidopsis, this tandem short stem-loop motif was found in *psbC* 3′ UTR, with good experimental support (AUROC=0.825), and in *ycf3* with relatively weaker support for its *in vivo* folding, given the higher reactivity within the second stem (Fig. 4e). Moreover, we identified several other shorter stem-loop motifs in the 3’ UTR of *ndhJ*, *rps18*, *rpl2*0, and *rps12*, where the *rpl2*0 and *rps12* are predicted with tandem stem-loops. Among these motifs, the tandem stem-loop in *rps12* showed good reactivity support (AUROC=0.80) (Supplementary Fig. 4f), in which the larger hairpin likely function as an IR motif. Altogether, the identified 3′ UTR structures demonstrate the evolutionary conservation of specific 3′ termini processing motifs, as well as indicate novel regulatory roles in transcript maturation, mRNA stability, and translation.

### Conservation of putative transcription terminator motif

Generally, plastid transcripts obtain a defined 3 ′ termini via post-transcriptional RNA processing (Zhelyazkova et al., 2012b), while transcription termination is commonly defined by stem-loop formation in nascent RNA in bacteria, such as rho-independent termination (Chen et al., 2019). We identified a conserved stem-loop motif that can be formed in both strands at the intergenic region between *petD* and *rpoA* operons expressed from the sense and antisense strand, respectively (Fig 4f-g). This motif, however, could not be validated using our DMS-MaP data. Nearly 40 years ago, a report on spinach plastid showed that the conserved stem-loop motif identified here might be important for transcription termination (Sijben-Müller et al., 1986), possibly helping avoid RNA polymerase collision during transcription of both strands, as shown for bacteria (Wang et al., 2023). However, transcriptional termination via RNA polymerase stalling by stem-loop structure in nascent transcript has not been shown for plastids yet.

### Conservation in 5**′** UTR motifs

The 5’ UTR in plastid mRNAs contains various *cis*-regulatory elements that typically interact with proteins to control translation, stability, and processing in response to developmental and environmental cues (Mayfield et al., 1994; Higgs et al., 1999; Zerges et al., 2003; Hirose and Sugiura, 2004). Here, we identified a conserved stem-loop (11-nt in length) in the *psbN* 5’ UTR (Fig. 5a), for which we are unaware of reported function. Unfortunately, chemical probing did not produce reliable mutation profile and experimental validation was, thus, unfeasible using our DMS-MaP dataset. Unlike most plastid 5’ UTRs, *psbN* 5’ UTR lacks a Shine-Dalgarno (SD)-like sequence and, instead, requires protein binding for translation initiation, e.g., binding to a 18-nt site adjacent to the start codon in tobacco *psbN* 5’ UTR (Kuroda and Sugiura, 2014). However, the identified conserved motif is often within 80-250nt (Supplementary Fig. 5a), hence, it is unlikely to play a direct role in translation initiation. Identification of primary and mature transcripts in barley revealed that the conserved motif is retained in the 5’ UTR of mature *psbN* transcript, but only in a subset of transcripts (Zhelyazkova et al., 2012a), suggesting a possible association between alternative 5’ processing and this conserved structure.

**Figure 5.**
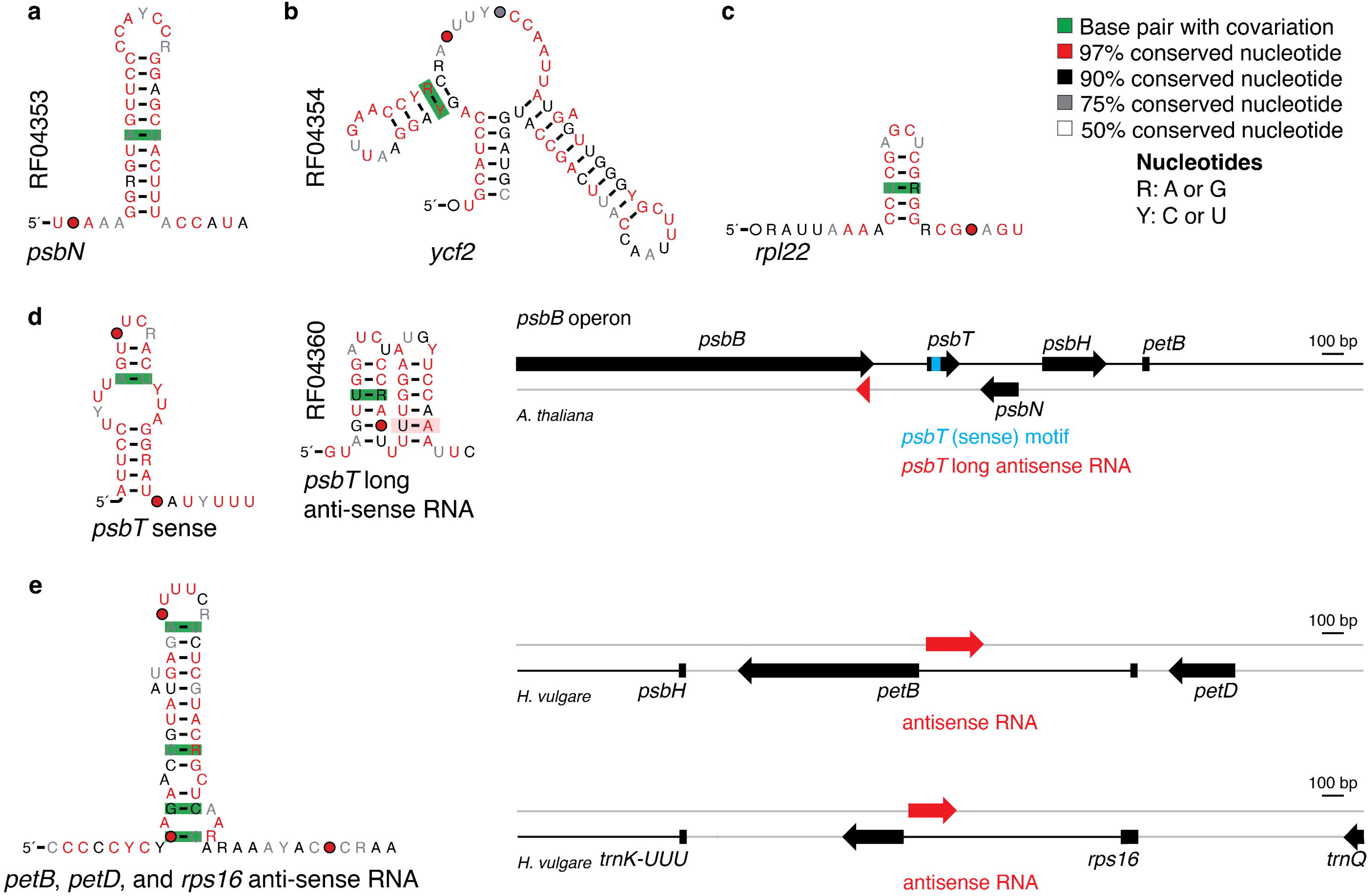
Conserved structure in 5’ untranslated region (UTR), coding sequence and antisense RNA. Consensus structure for motif identified in the 5’ UTR of *psbN* **(a)** and *ycf2* **(b)**, coding sequence of *rpl22* **(c)**. **(d)** Consensus structure for motif identified in *psbT* coding sequence (*psbT* sense, left) and in *psbT* long antisense RNA (middle). Genomic context of the *psbT* sense RNA motif (blue) and antisense RNA motif (red) in *Arabidopsis thaliana* plastid (right). **(e)** Consensus structure of antisense RNA motif to *petB*, *petD*, and *rps16* (left). Genomic context of the antisense RNA (red) in *Hordeum vulgare* mapping to group II intron of *petB* and *rps16* (right). **(a-b, d)** Rfam accessions provided on the left side of the structures (vertical text).

Many plastid genomes contain two inverted repeats (IRa and IRb) that separate a large single copy region (LSC) from a short single copy region (SSC), hence, genes in inverted repeat are typically found duplicated in the genome. We identified a conserved motif in the 5’ UTR of both copies of *ycf2* (Fig. 5b). This motif, however, is associated with a tRNA in the opposite DNA strand, and selective pressure is likely on the tRNA structure, rather than on *ycf2* 5’ UTR. Although such motifs are usually discarded from RNA conservation analyses, it is possible that tRNA-like structure in the 5’ UTR has regulatory function, and the tRNA-like and *ycf2* 5’ UTR association has been retained in both gene copies. For instance, the *psbDCZ* operon has a similar tRNA-like structure between *psbC* and *psbZ*, which has been proposed to function in polycistronic mRNA maturation via tRNA-induced RNase cleavage, yielding mono-cistronic species (Gawroński et al., 2020). Similarly, in mitochondria, tRNA-induced RNase cleavage has been shown to regulate maturation of transcripts containing tRNA-like structure (Okui et al., 2016). It is possible that the tRNA-like motif identified in *ycf2* 5’ UTR is required in mRNA maturation to define precise 5’ end; however, this requires further experimental evaluation. In addition, we identified novel conserved motifs in the 5’ UTR of a protease complex subunit (*clpP*) and a chloroplast-encoded RNA polymerase subunit (*rpoB*) (Supplementary Fig. 5b), for which we are also unaware of described function. Altogether, we have identified three novel conserved motifs located in the 5’ UTR of diverse transcripts, likely involved in post-transcriptional regulation.

### Conservation in motifs within coding region and antisense noncoding RNA

The large ribosomal subunit *rpl22* is essential in most seed plants (Fleischmann et al., 2011). Here, we identified a conserved motif in the *rpl22* coding sequence (CDS), formed by a short stem-loop, likely uncharacterized for its function in plastids (Fig. 5c), as well as a similar stem-loop identified in the *rpl2* CDS (Supplementary Fig. 5c). *rpl22* and *rpl2* genes are often part of a large operon with other ribosomal protein genes (e.g., *rps19*, *rpl2*3, and rps3), for which maturation requires activity of endonuclease at multiple sites (Daniell et al., 2016), while splicing and editing seem either less prevalent or non-existing in *rpl2* and *rpl22* across plant species. Given the tight regulation of ribosomal subunits at multiple levels, it is possible that the conserved motif within *rpl2* and *rpl22* CDS plays a regulatory role possibly by interacting with specific proteins.

We identified a novel conserved motif in the *psaA* CDS, near the 3’ (Supplementary Fig. 5c), and another in *psbT* CDS (Fig. 5d), both of which yet uncharacterized for their function. *psbT* is part of the *psbB* operon, which has *psbN* on the opposite genomic strand situated between *psbT* and *psbH* genes. It has been shown that *psbT* expression is controlled by two antisense RNAs originated from *psbN* transcription by the nuclear-encoded polymerase (NEP), in which the short antisense RNA hybridizes to *psbT* CDS, thereby stabilizing *psbT* transcript while blocking its translation during photooxidative stress (Zghidi-Abouzid et al., 2011). It is thus possible that the conserved motif in *psbT* CDS is involved in the regulation of sense-antisense interaction with the *psbN*-derived noncoding RNA. Interestingly, we identified a conserved motif mapping to the long, but not the short antisense RNA derived from *psbN* transcription (Fig. 5d), suggesting a complex and fine-tuned regulation of *psbT* and, potentially the entire *psbB* operon, involving RNA motifs conserved in seed plants, fern, moss, and green algae (Supplementary Fig. 5d).

We identified another conserved motif antisense to a coding gene; however, in the opposite strand to group II intron D5 and D6 in *petB*, *petD*, *rps16* (Fig. 5e). In barley, RNA was also found in the antisense region of *petB* intron (Zhelyazkova et al., 2012a), suggesting a regulatory function for conserved motif in such noncoding antisense RNAs. Furthermore, beyond the conserved motifs associated with genic context, we identified eight conserved motifs likely associated with noncoding RNA for which evidence for transcription is unclear (Supplementary Fig. 6a-b). For instance, DMS reactivity supported the *in vivo* folding of one of the identified non-coding motifs (Supplementary Fig. 6b), demonstrating that the RNA is transcribed and potentially functional in plastids.

## CONCLUSIONS

This study presents a systematic identification of conserved RNA structures in plastids across the plant phylogenetic tree, enabled by the development of a novel genome-wide pipeline for the identification of conserved structures across thousands of plastid genomes. We validated our discovery pipeline by (1) confirming recovery of various well-characterized RNA structures (e.g., rRNA, tRNAs, group II intron domain, and 3’ end processing stem-loops), (2) experimental validation of *in vivo* RNA folding in plastid, and (3) cross-validation of structural models by Rfam. In addition, we discovered novel and uncharacterized motifs, such as those within the 5’ UTR of *psbN* (also conserved in cyanobacteria), coding sequence of *psbT* and *rpl22*, intronic regions of *atpF*, *rpl16*, *ycf3*, and *clpP*, and antisense RNAs to *psbT*, *petB*, *petD*, and *rps16*. During curation, the Rfam team performed a species distribution analysis across all domains of life. This initial broader phylogenetic analysis revealed that the motif associated with *psbI*/*ycf3*/*rps4*/*chlN* 3′ termini (RF04351) and *psbN* 5′

UTR (RF04353) are conserved in cyanobacteria, suggesting shared regulatory functions with plastid ancestral. Altogether, our bioinformatic approach identified a large set of novel conserved RNA structures in plastids, cross-validated and available in Rfam.

These findings significantly expand the known repertoire of putative *cis*- and *trans*-regulatory RNA elements in plants, evidencing that conserved RNA structure is a pervasive feature of plastid gene regulation. Our work will be the foundation for the discovery of novel regulatory mechanisms in plastids.

## METHODS

### Plastid genomic sequences

An initial list of 3823 plastid genomes from seed plants (i.e., spermatophyta) was obtained from the CpGDB database (Singh et al., 2020). This list was complemented with plastid genomes obtained directly from NCBI (July 11, 2024) representing all available photosynthetic organisms, resulting in a total of 14034 plastid genomes.

### RNA structure search using Rfam database

Covariance models (.cm) of 4178 known RNA families were obtained from Rfam database (v15.0) (Ontiveros-Palacios et al., 2025) and cmsearch (from Infernal v1.1.4 package) (Nawrocki and Eddy, 2013) with --ga threshold was performed against 14034 plastid genome sequences and any hits found were considered as RNA structure homologs.

### Genome-wide identification of conserved RNA structure

The pipeline developed in this work is available from GitHub (https://github.com/RodrigoReisLab/RNA_conservation) and as a container at Docker Hub Container Image Library (https://hub.docker.com/r/dollycm/rnatools). First, we produced sequence windows of 250 nt with an overlap of 75 nt covering 3823 plastid genomic sequence (seed plants), totalling ∼5.6M windows. Windows were processed to remove sequences comprised of a single nucleotide type (e.g., A, T, G, or C), containing regions with unsequenced nucleotides (i.e., marked as ‘N’), or redundant sequences (i.e., 100% identical). Sequences were then clustered using MMSeqs2 package (easy-cluster) (Steinegger and Söding, 2017) with 95% sequence identity to identify highly similar sequences, which were then removed with only one representative sequence retained per cluster. Representative sequences were further clustered with a minimum of 50% sequence identity, resulting in a total of 10431 clusters.

Clustered sequences were aligned with Clustal Omega (Sievers and Higgins, 2021) and RNALalifold (Lorenz et al., 2011) was used to predict folding for each sequence, which was performed with temperature set to 21°C / 294.15K (-T option) and no lone pairs (--noLP) to identify sequences in a cluster that share a minimum of 2 base pairs. Approximately 30% of the clusters (3136/10431) passed this screening step. A progressive RNA structure prediction was performed using LocARNA package (mlocarna --probabilistic) (Will et al., 2012) on the unaligned sequences obtained from MMSeqs2 with parameters set to --local-progressive and --rnafold-temperature=21°C. We identified 1827 clusters with structures sharing a minimum of 3 base pairs.

Alignments produced with LocARNA had a predefined length of 250 nt (i.e., input window size), hence, they were trimmed to only include the identified motif and additional 20 nt in each extremity using an in-house PERL program. Trimmed alignments were evaluated using our in-house script, to only retain sequences with at least 75% of base pairs in each stem, at least 50% of entire motif in each stem, and at least 30% of GC content and removed any redundant sequences. This would ensure that each stem-loop is formed independently with a defined 30% GC content, providing sufficient flexibility to accommodate different conformations while avoiding overly rigid or overly flexible stems while allowing for structural variations (like 3-stem, 4-stem, 5-stem tRNAs) to be retained. Only clusters with at least 7 sequences in the alignment, were considered further. Cleaned alignments were converted into covariance models using cmbuild and cmcalibrate (Infernal v1.1.4) (Nawrocki and Eddy, 2013) and searched against the 3823 plastid genomes from seed plants to find homologs with cmsearch and cmalign programs of Infernal v1.1.4. Output of Infernal were processed using previously described PERL scripts to reformat Infernal output (Barrick, 2009). Sequences were re-evaluated with same criteria as above, except for 15% GC content (instead of 30%)to obtain non-redundant homologs that adopt the predicted structure and produce a new alignment that could be used as seed. Only clusters with atleast 7 sequences that were capable of finding homologs were considered further. Covariation analysis was performed using R-scape (Rivas et al., 2016) with two-tailed test and --cacofold to identify statistically significant covarying base-pairs, indicative of conservation of structure beyond phylogeny, and any additional base-pairing missed out by LocARNA. Only the clusters with at least 2% covariation were considered as reliable alignments. The structures obtained with LocARNA and R-scape --cacofold were compared qualitatively and, if the predicted structures were completely different, the LocARNA predicted structures were considered, since it is likely to be the more biologically relevant structure. To further check the sequence-structure compatibility, nhmmer (Wheeler and Eddy, 2013) was used with default parameters, for which the clustered sequences were de-aligned and are re-aligned independently without structure information to correct for sequence misalignment, which may arise due to structure positions bias during LocARNA simultaneous fold and align. Structure consensus predicted with LocARNA were then added to nhmmer alignments and subjected to evaluation with R-scape (--cacofold), i.e., only alignments for which predicted structures with LocARNA and CaCoFold showed consistent overall architecture and at least one base-pair with statistically significant covariation were considered reliable and high confidence predictions. This approach resulted in the discovery of 57 conserved motifs in plastid genomes. CaCoFold alignments were then used as seed for each conserved motifs to obtain covariance models built with cmbuild and cmcalibrate programs of Infernal (Nawrocki and Eddy, 2013). Covariance models were used against all 14034 plastid genomes with e-value 0.01 to search for homologs using cmsearch program of Infernal v1.1.4. The obtained hits were processed using the in-house script with same parameters as above to evaluate each identified homolog. High confidence RNA structures post context analysis were independently assessed by the Rfam team and deposited in Rfam database.

Our pipeline can be readily applied to any genome or set of sequences, including selective usage at any analysis step, composable, and adaptable to user needs.

### Genomic context and phylogenetic analyses of conserved RNA structures

To identify the genomic context, RNA coordinates were mapped to gene annotation (*.gff3*) for the corresponding plastid genome using bedtools (Quinlan and Hall, 2010) closest with options -s (gene and the RNA are on the same strand) -D b (reports if genes are upstream or downstream to the RNAs). bedtools outputs were analysed using in-house PERL and BASH scripts and only RNAs within 500 nt of a gene were considered as homologs and further used to categorize RNA motifs. We employed the following broad categories for genome context: tRNA, 5’ UTR, 3’ UTR, and exon-intron junction. Additional curation was carried out for tRNA motif homologs with bitscore threshold used was half of the highest scoring RNA homolog and with a distance of 0 (overlapping with genes). If multiple motif map to the same RNA, the highest scoring RNA is retained as a homolog. To obtain the phylogenetic distribution of each conserved RNA motif, NCBI taxonomic information (taxon ID and taxonomic lineage) were obtained using edirect utilities (Kans, 2025) from provided NCBI genome IDs. The mapping was obtained for genera across 586 plant families from which a dataset compatible with iTOL (Letunic and Bork, 2024) was generated containing percentage of genera within a plant family carrying RNA motifs. The dataset was mapped to the NCBI *Viridiplantae* tree in iTOL (https://itol.embl.de/) for visualization.

### RNA DMS probing *in vivo*

*In vivo* DMS probing was performed as previously described (Ding et al., 2015) with minor modifications. Three biological replicates of five-day-old *Arabidopsis thaliana* Col-0 seedlings grown in liquid half-strength Murashige and Skoog medium supplemented with 0.25% (w/v) sucrose were immersed in DMS reaction buffer (40 mM HEPES-KOH, pH 7.5, 0.5 mM MgCl₂, and 100 mM KCl). For DMS-treated samples, DMS was added to a final concentration of 1% (v/v), whereas control samples underwent the same procedure without DMS treatment. Samples were vacuum infiltrated and incubated at room temperature for a total of 15 min. The probing reaction was quenched by adding DTT to a final concentration of 0.5 M, followed by vigorous vortexing until the DTT was completely dissolved. The seedlings were washed twice with RNase-free water, immediately frozen in liquid nitrogen, and total RNA was extracted using the Quick-RNA MiniPrep Kit (Zymo Research).

### DMS mutation profiling (MaP) library preparation

DMS-MaP library preparation was performed as previously described (Mitchell et al., 2023) with minor modifications. Poly(A)-enriched RNA was fragmented in NEBNext First Strand Synthesis Reaction Buffer (NEB) at 94 °C for 4 min, followed by purification using Zymo RCC-5 column (Zymo Research) and elution in 6.5 μL of RNase-free water. For reverse transcription, 200 ng random nonamers were added to the fragmented RNA, and the mixture was incubated at 68°C for 5 min followed by cooling to room temperature to allow primer annealing. First-strand cDNA synthesis was performed with 0.5 μL ultraMarathonRT (RNAConnect; R1002M) in a 20-μL DMS-MaP reaction containing 50 mM Tris-HCl (pH 8.3), 200 mM KCl, 5 mM DTT, 0.5 mM dNTPs, 1 mM MnCl_2_, 0.4 U/μL RNase inhibitor, and 20% (v/v) glycerol. Reverse transcription was carried out at 42°C for 3 h, followed by enzyme inactivation at 95°C for 3 min and immediate cooling on ice. Subsequently, 1 μL of 20 mM EDTA was added to terminate the reaction, and the mixture was incubated at room temperature for 5 min. MgCl_2_ was then added to a final concentration of 6 mM. Second stranded cDNA synthesis and all subsequent library preparation steps were performed using the NEBNext Ultra II DNA Library Prep Kit for Illumina (NEB) according to the manufacturer’s instructions. cDNA libraries were sequenced by Novogene Co. (Munich, Germany) for 150-bp paired-end Illumina sequencing.

### DMS mutation profiling (MaP) data analysis

Illumina paired-end sequencing data (fastq) were processed using nfcore/rnaseq v3.12.0 with default parameters to obtain trimmed reads. Replicates were initially merged and aligned to *Arabidopsis thaliana* chloroplast genome (NC_000932.1) using star-salmon (*-- alignIntronMax 5000, --seqBias and --gcBias*) to assess gene coverage. Genomic sequences encompassing conserved RNA structure, including 150-nt flanking sequences, were used as reference for mapping using the software shapemapper2 (modular approach) (Shen et al., 2018). We then used RNA Framework (Incarnato et al., 2018) for mutation count and normalization. Mutations were counted in mapped reads (BAM files) with rf-count (*-ds 100 -m -ncl -es -ni -na -me 0.1 -dc 3 -mm*) and adenine and cytosine counts were extracted with rf-mmtools (*-mpr 2 -kb AC*). Mutation counts were normalized with rf-norm (*-sm 4 -nm 3 -rb AC -ec 300)* with normalization factor obtained with rf-normfactor. Finally, we used rf-eval to evaluate the agreement between predicted structures and *in vivo* RNA probing data (AUROC ≥ 0.50). Structures were visualized displaying RNA probing reactivities with RfviewJS (https://github.com/dincarnato/RFviewJS/).

## DATA AVAILABILITY

DMS-MaP data is available from NCBI SRA (BioProject PRJNA1514713). The pipeline developed in this work is available at GitHub (github.com/RodrigoReisLab/RNA_conservation) and the tools used in the pipeline are available as a container at Docker Hub Container Image Library (https://hub.docker.com/r/dollycm/rnatools). All novel conserved RNA motifs have been submitted to Rfam and RNA Stockholm alignments are available at Bitbucket (https://bitbucket.org/rodrigoreislab/chloroplast_motif_families).

## ACKNOWLEDGEMENTS

The authors thank the Rfam team members Blake Sweeney and Nancy Ontiveros for helpful discussions and curation of our identified motifs. We thank Elena Rivas (Harvard University) for valuable discussions and feedback. We thank Danny Incarnato (University of Groningen) for assisting with RNA Framework implementation. We also thank Michael Crawford and Rachappa Balkunde (Bayer Crop Science) for guidance on applicability for crop protection. This work was supported by the Swiss National Science Foundation (PCEFP3_203328 and 320030-231536) and the Grants4Ag Program (Bayer Crop Science).

## AUTHOR CONTRIBUTIONS

RSR and DM conceived and designed this work. DM performed all bioinformatic analyses. CX performed DMS probing and library preparation. JH contributed to initial experiments and planning. RSR and DM wrote the manuscript. All authors read and approved the manuscript.

## SUPPLEMENTARY INFORMATION

**Supplementary Table 1**. Rfam motifs identified in plastid genomes. List of Rfam IDs identified in the corresponding number of chloroplast genomes. IDs marked with asterisk indicate predicted structure in plastid genomes for which we could not identify previously described function in plastids.

**Supplementary Table 2**. Conserved RNA structures identified in this study.

**Supplementary Table 3**. DMS probing data from *Arabidopsis thaliana*.

**Supplementary Fig. 1.** Consensus structure models for tRNA and stem-loop structure similar to bacterial 16S rRNA helix 8. **(a)** tRNA models. (b) Stem-loop structure similar to bacterial 16S rRNA helix 8. **(c)** DMS reactivity for tRNA structures identified in *Arabidopsis thaliana*; value <0.3 (black), 0.3-0.7 (yellow), and >0.7 (red) indicate low, medium, and high reactivity (i.e., unpairness), respectively.

**Supplementary Fig. 2.** Associated genes and consensus structure models of group II intron D5. **(a)** Distribution of genes identified with group II intron D5 from different models. Inset shows the distribution of genes with less than 50 representatives. **(b)** Consensus structure of five models representing group II intron D5 mapped to *ndhA*, *ndhB*, *petB*, *atpF*, *ycf3*, and *rps12* intron. **(c)** Consensus structure of model representing group II intron D5 mapped to *trnK* intron. **(d)** DMS reactivity for structures identified in *Arabidopsis thaliana*; value <0.3 (black), 0.3-0.7 (yellow), and >0.7 (red) indicate low, medium, and high reactivity (i.e., unpairness), respectively. Area under the receiver operating characteristic curve (AUROC; 0 to 1) indicates the agreement between structure and reactivity profile, in which high AUROC values correspond to better agreement.

**Supplementary Fig. 3.** Associated genes and consensus structure models of group II intron D1 and D3. **(a)** Consensus structure of models representing group II intron D1 and D3 mapped to *ClpP* and *trnK* intron, respectively. **(b)** DMS reactivity for structures identified in *Arabidopsis thaliana*; value <0.3 (black), 0.3-0.7 (yellow), and >0.7 (red) indicate low, medium, and high reactivity (i.e., unpairness), respectively. Area under the receiver operating characteristic curve (AUROC; 0 to 1) indicates the agreement between structure and reactivity profile, in which high AUROC values correspond to better agreement.

**Supplementary Fig. 4.** Consensus structure models and genome distribution of motifs in 3′ UTR. Consensus structure of models representing 3′-termini processing motifs in *psbA* **(a),** *ccsA*-*ndhD* **(b),** *psbJ* **(c). (d)** Genomes identified with different models of 3 ′ -termini processing motif. All three model1 find homologs in most plastid genomes, hence general models, while the other models are more genome specific and complement homolog identification by model1 **(e)** Consensus structure of motifs identified in 3′UTR with unknown function, found associated with *ndhJ*, *rpl2*0, *rps18* and *rps12*. **(f)** DMS reactivity for structures identified in *Arabidopsis thaliana*; value <0.3 (black), 0.3-0.7 (yellow), and >0.7 (red) indicate low, medium, and high reactivity (i.e., unpairness), respectively.

**Supplementary Fig. 5.** Distribution and models for 5′UTR, coding sequence, and antisense RNAs. **(a)** Histogram shows the distance distribution of motif (consensus structure in Fig 5a) in the 5′UTR of *psbN* gene. Consensus structure of motifs found in **(b)** 5′UTR of *clpP* and *rpoB* **(c)** coding sequence of *rpl2* and *psaA*-psaB. **(d)** Phylogenetic distribution of novel conserved RNA motifs in *psbT* coding sequence and anti-sense RNA to *psbT* and *petB* Circle size (outer layer) represents the percentage distribution of RNA motif in given plant clade (different color branches).

**Supplementary Fig. 6.** Consensus structure of motifs with no clear genic context. **(a)** Consensus structures. **(b)** Consensus structures and corresponding DMS reactivity for structure identified in *Arabidopsis thaliana*; value <0.3 (black), 0.3-0.7 (yellow), and >0.7 (red) indicate low, medium, and high reactivity (i.e., unpairness), respectively.

