## Supplementary Figure for "RNA structure conservation in plastids across plant evolution"

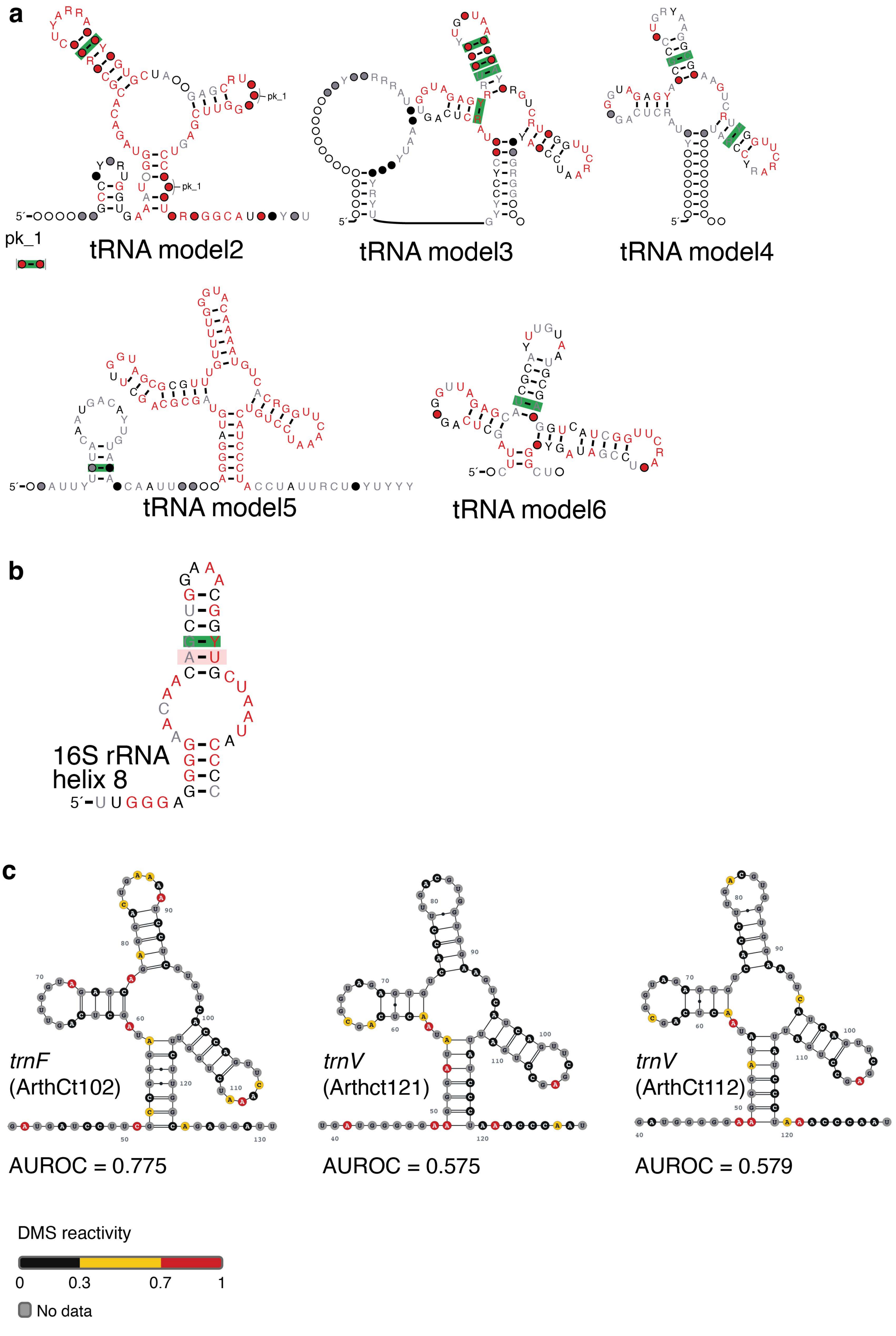


**Supplementary Fig. 1**. Consensus structure models for tRNA and stem-loop structure similar to bacterial 16S rRNA helix 8. (a) tRNA models. (b) Stem-loop structure similar to bacterial 16S rRNA helix 8. (c) DMS reactivity for tRNA structures identified in Arabidopsis thaliana; value <0.3 (black), 0.3-0.7 (yellow), and >0.7 (red) indicate low, medium, and high reactivity (i.e., unpairness), respectively.

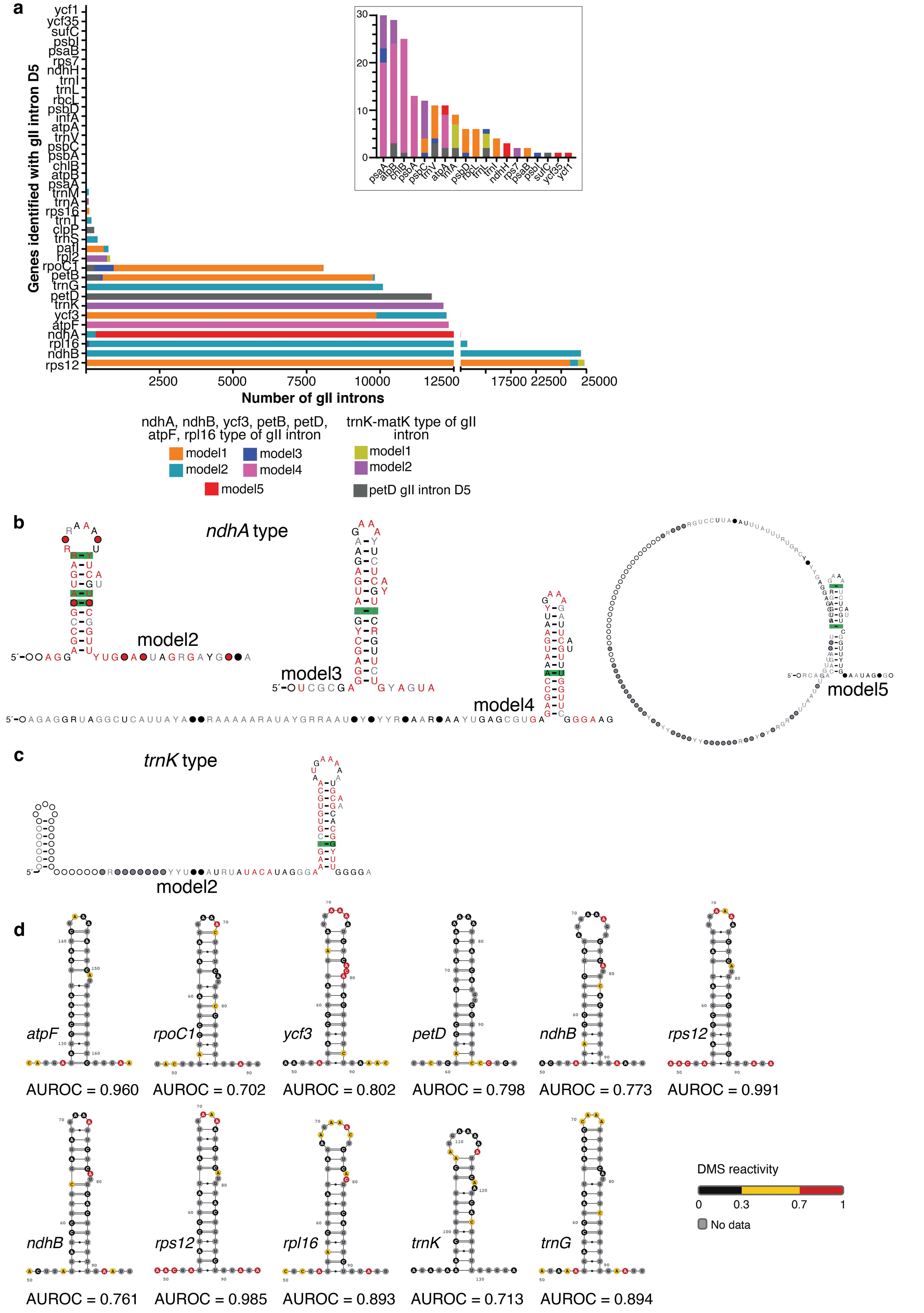


**Supplementary Fig. 2**. Associated genes and consensus structure models of group II introns. (a) Distribution of genes identified with group II intron D5 from different models. Inset shows the distribution of genes with less than 50 representatives. (b) Consensus structure of five models representing group II intron D5 mapped to ndhA, ndhB, petB, atpF, ycf3, and rps12 intron. (c) Consensus structure of model representing group II intron D5 mapped to trnK intron. (d) DMS reactivity for structures identified in Arabidopsis thaliana; value <0.3 (black), 0.3-0.7 (yellow), and >0.7 (red) indicate low, medium, and high reactivity (i.e., unpairness), respectively. Area under the receiver operating characteristic curve (AUROC; 0 to 1) indicates the agreement between structure and reactivity profile, in which high AUROC values correspond to better agreement.

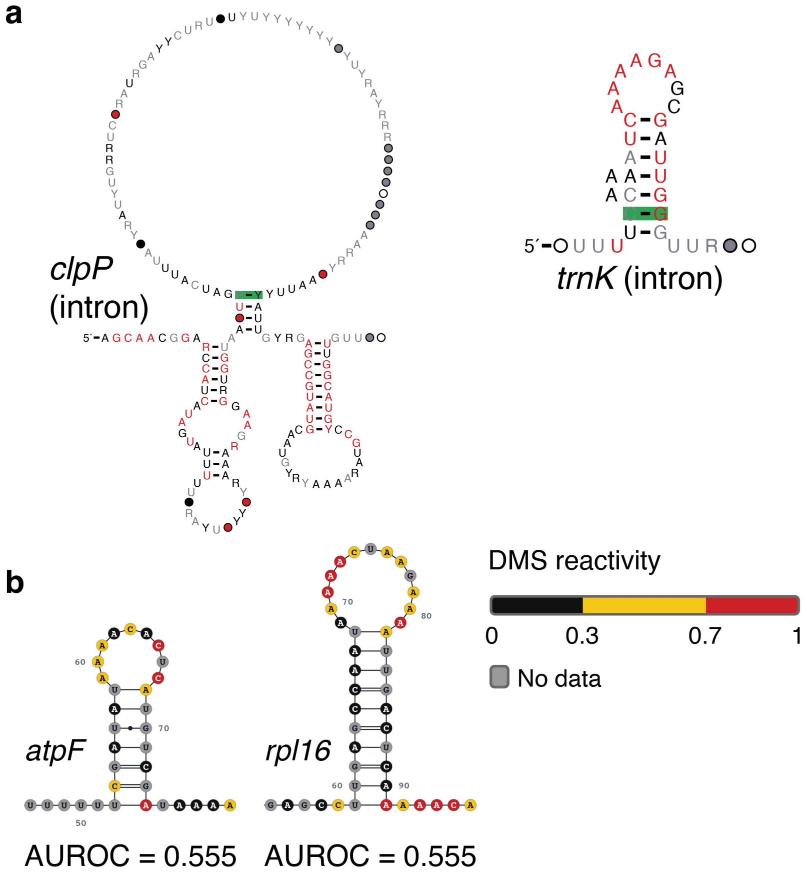


**Supplementary Fig. 3**. Associated genes and consensus structure models of group II intron D1 and D3. (a) Consensus structure of models representing group II intron D1 and D3 mapped to ClpP and trnK intron, respectively. (b) DMS reactivity for structures identified in Arabidopsis thaliana; value <0.3 (black), 0.3-0.7 (yellow), and >0.7 (red) indicate low, medium, and high reactivity (i.e., unpairness), respectively. Area under the receiver operating characteristic curve (AUROC; 0 to 1) indicates the agreement between structure and reactivity profile, in which high AUROC values correspond to better agreement.

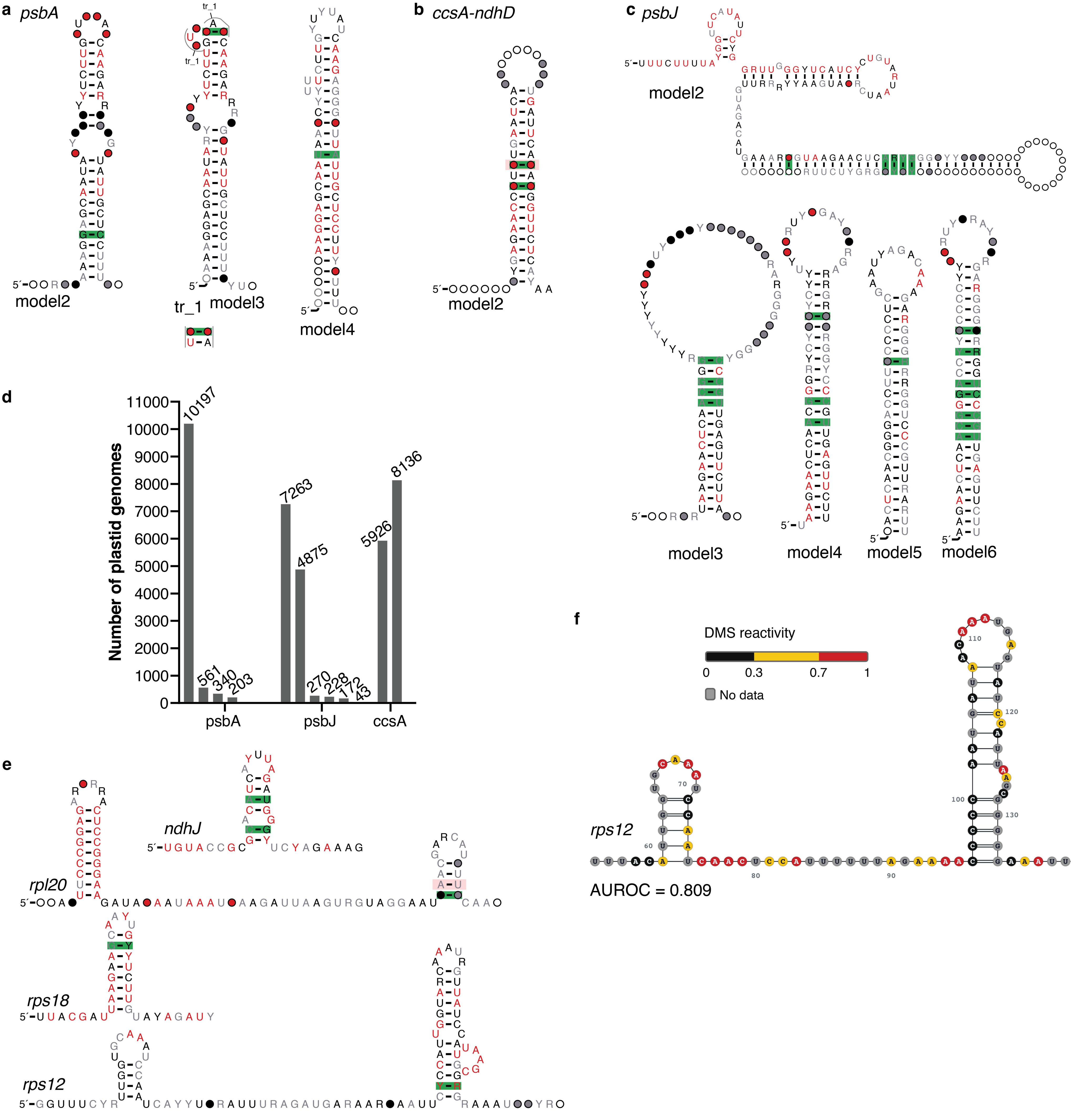


**Supplementary Fig. 4**. Consensus structure models and genome distribution of motifs in 3′ UTR. Consensus structure of models representing 3′-termini processing motifs in psbA (a), ccsA-ndhD (b), psbJ (c). (d) Genomes identified with different models of 3′-termini processing motif. All three model1 find homologs in most plastid genomes, hence general models, while the other models are more genome specific and complement homolog identification by model1 (e) Consensus structure of motifs identified in 3′UTR with unknown function, found associated with ndhJ, rpl20, rps18 and rps12. (f) DMS reactivity for structures identified in Arabidopsis thaliana; value <0.3 (black), 0.3-0.7 (yellow), and >0.7 (red) indicate low, medium, and high reactivity (i.e., unpairness), respectively.

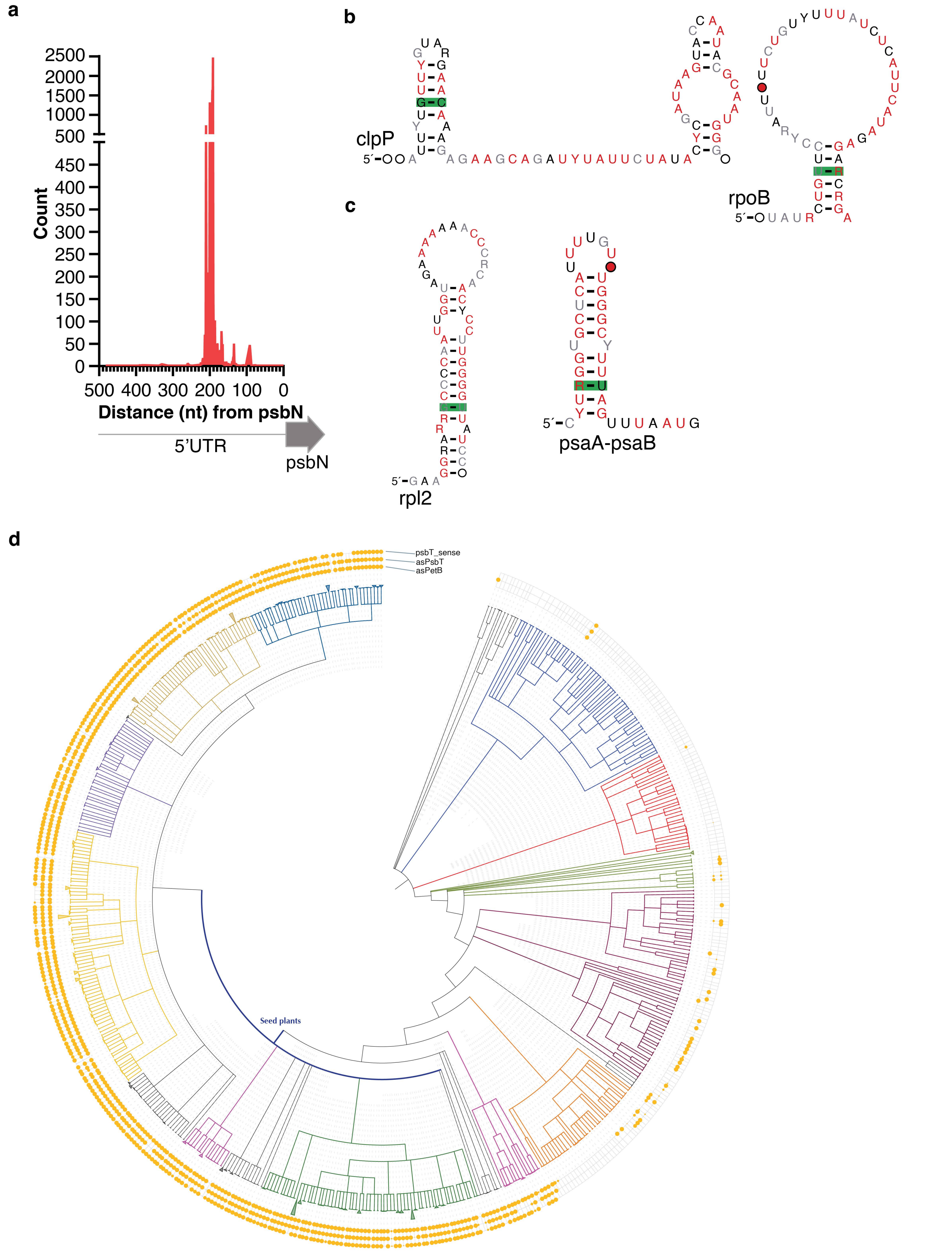


**Supplementary Fig. 5**. Distribution and models for 5′UTR, coding sequence, and antisense RNAs. (a) Histogram shows the distance distribution of motif (consensus structure in Fig 5a) in the 5′UTR of psbN gene. Consensus structure of motifs found in (b) 5′UTR of clpP and rpoB (c) coding sequence of rpl2 and psaA-psaB. (d) Phylogenetic distribution of novel conserved RNA motifs in psbT coding sequence and anti-sense RNA to psbT and petB Circle size (outer layer) represents the percentage distribution of RNA motif in given plant clade (different color branches).

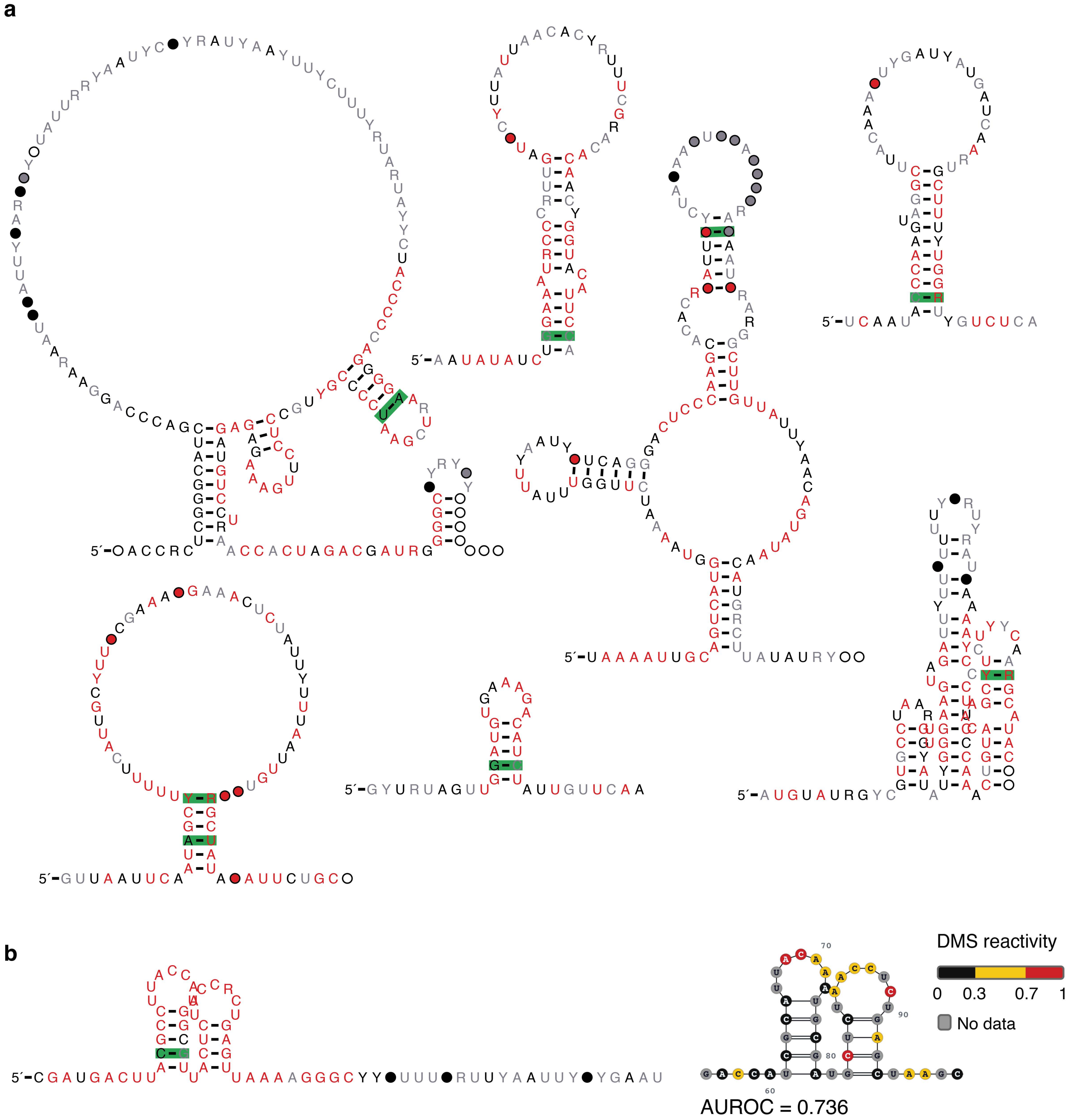


**Supplementary Fig. 6**. Consensus structure of motifs with no clear genic context. (a) Consensus structures. (b) Consensus structures and corresponding DMS reactivity for structure identified in Arabidopsis thaliana; value <0.3 (black), 0.3-0.7 (yellow), and >0.7 (red) indicate low, medium, and high reactivity (i.e., unpairness), respectively.
