## Supplementary Table 1 for "RNA structure conservation in plastids across plant evolution"

**Supplementary Table 1**. Rfam motifs identified in plastid genomes. List of Rfam IDs identified in the corresponding number of chloroplast genomes. IDs marked with asterisk indicate predicted structure in plastid genomes for which we could not identify previously described function in plastids.

| **Rfam ID** | **description of the Rfam motif** | **Number of Plastid Genomes** |
| --- | --- | --- |
| RF00177 | SSU_rRNA_bacteria | 14029 |
| RF00005 | tRNA | 14029 |
| RF01959 | SSU_rRNA_archaea | 14027 |
| RF02542 | SSU_rRNA_microsporidia | 14021 |
| RF02541 | LSU_rRNA_bacteria | 14019 |
| RF01960 | SSU_rRNA_eukarya | 14017 |
| RF02540 | LSU_rRNA_archaea | 13951 |
| RF01419* | isrR-Antisense RNA which regulates isiA expression | 13948 |
| RF00001 | 5S_rRNA | 13900 |
| RF00029 | Intron_gpII | 13792 |
| RF00028 | Intron_gpI | 13497 |
| RF01852 | tRNA-Sec | 7320 |
| RF02004 | group-II-D1D4-5 | 27 |
| RF00002 | 5_8S_rRNA | 8 |
| RF02033 | HEARO-HNH endonuclease-associated RNA and ORF (HEARO) RNA | 2 |
| RF03545 | Flavi_ISFV_CRE-Insect-specific Flavivirus 3' UTR cis-acting replication element (CRE) | 1 |
| RF02711* | TeloSII_ncR49 | 167 |
| RF00010* | RNaseP_bact_a | 145 |
| RF00169* | Bacteria_small_SRP | 143 |
| RF00023* | tmRNA | 127 |
